# Antipsychotic behavioral phenotypes in the mouse Collaborative Cross recombinant inbred inter-crosses (RIX)

**DOI:** 10.1101/761353

**Authors:** Paola Giusti-Rodríguez, James G. Xenakis, James J. Crowley, Randal J. Nonneman, Daniela M. DeCristo, Allison Ryan, Corey R. Quackenbush, Darla R. Miller, Ginger D. Shaw, Vasyl Zhabotynsky, Patrick F. Sullivan, Fernando Pardo-Manuel de Villena, Fei Zou

**Affiliations:** Department of Genetics, University of North Carolina, Chapel Hill, NC, USA; Department of Biostatistics, University of North Carolina, Chapel Hill, NC, USA; Department of Psychiatry, University of North Carolina, Chapel Hill, NC, USA; Department of Medical Epidemiology and Biostatistics, Karolinska Institutet, Stockholm, Sweden; Lineberger Comprehensive Cancer Center, University of North Carolina, Chapel Hill, NC, USA

## Abstract

Schizophrenia is an idiopathic disorder that affects approximately 1% of the human population, and presents with persistent delusions, hallucinations, and disorganized behaviors. Antipsychotics are the standard pharmacological treatment for schizophrenia, but are frequently discontinued by patients due to inefficacy and/or side effects. Chronic treatment with the typical antipsychotic haloperidol causes tardive dyskinesia (TD), which manifests as involuntary and often irreversible orofacial movements in around 30% of patients. Mice treated with haloperidol develop many of the features of TD, including jaw tremors, tongue protrusions, and vacuous chewing movements (VCMs). In this study, we used genetically diverse Collaborative Cross (CC) recombinant inbred inter-cross (RIX) mice to elucidate the genetic basis of antipsychotic-induced adverse drug reactions (ADRs). We performed a battery of behavioral tests in 840 mice from 73 RIX lines (derived from 62 CC strains) treated with haloperidol or placebo in order to monitor the development of ADRs. We used linear mixed models to test for strain and treatment effects. We observed highly significant strain effects for almost all behavioral measurements investigated (p<0.001). Further, we observed strong strain-by-treatment interactions for most phenotypes, particularly for changes in distance traveled, vertical activity, and extrapyramidal symptoms (EPS). Estimates of overall heritability ranged from 0.21 (change in body weight) to 0.4 (VCMs and change in distance traveled) while the portion attributable to the interactions of treatment and strain ranged from 0.01 (for change in body weight) to 0.15 (for change in EPS). Interestingly, close to 30% of RIX mice exhibited VCMs, a sensitivity to haloperidol exposure, approximately similar to the rate of TD in humans chronically exposed to haloperidol. Understanding the genetic basis for the susceptibility to antipsychotic ADRs may be possible in mouse, and extrapolation to humans could lead to safer therapeutic approaches for schizophrenia.

## INTRODUCTION

Schizophrenia is a chronic, severe, and disabling brain disorder that affects about 1% of the population worldwide and is associated with substantial morbidity, mortality, and personal and societal costs (Saha *et al*. 2007; World Health Organization 2008; Laursen *et al*. 2012; Cloutier *et al*. 2016). Onset typically occurs in adolescence or early adulthood with the emergence of a number of symptoms, including hallucinations, delusions, and disorganized behavior (American Psychiatric Association. 2013). Antipsychotic medications are the mainstay of treatment for schizophrenia, but as much as 75% of patients discontinue assigned treatments due to side effects and/or inefficacy over relatively short periods of time (Lieberman *et al*. 2005). Deciphering the pharmacogenetics of antipsychotics could eventually allow the development of predictive algorithms, such that physicians could predict which patients are prone to develop side effects or less likely to achieve a therapeutic response. Advances in this arena could lead to safer and more efficacious therapeutic interventions for the treatment of schizophrenia.

Haloperidol is a high-potency, typical, first-generation antipsychotic whose use is associated with motoric adverse drug reactions (ADRs), including tardive dyskinesia (Soares-Weiser and Fernandez 2007), involuntary orofacial movements, and extrapyramidal symptoms (i.e., dystonia and Parkinsonism) (Miller *et al*. 2008). There is substantial inter-individual variation in liability to these ADRs and direct and indirect evidence suggest a role for genetic variation (Lerer *et al*. 2005; Patsopoulos *et al*. 2005; Bakker *et al*. 2006). There is significant heterogeneity in therapeutic response to antipsychotics (Lieberman *et al*. 2005), with roughly equal proportions of patients experiencing remission, partial response, and no benefit. Vacuous chewing movements (VCMs), which present in rodents as purposeless mouth openings in the vertical plane, are a valid rodent model for the human pharmacogenetic phenotype of tardive dyskinesia (Tomiyama *et al*. 2001; Turrone *et al*. 2002a; Crowley *et al*. 2012b; Crowley *et al*. 2014). Indeed, using 27 genetically diverse inbred mouse strains, we previously showed that chronic haloperidol treatment (60 days) can bring on VCMs that persist for 1.5 years, that VCMs and extrapyramidal side effects are highly heritable (∼0.9), and that there are strong strain effects (Crowley *et al*. 2012a). Genome-wide association mapping of these same 27 inbred strains nominated ∼50 genes for association with haloperidol-induced ADRs (Crowley *et al*. 2012b). Furthermore, using RNA sequencing of striatal tissue from mice chronically treated with haloperidol, we previously observed an overlap between the genetic variation underlying the pathophysiology of schizophrenia and the molecular effects of haloperidol (Kim *et al*. 2018).

Traditional recombinant inbred strains that are generated by crossing two parental inbred strains followed by 20 generations of inbreeding by brother-sister mating lack genetic diversity and are limited by the number of available strains (e.g. C57BL6/J, 129S1, BALB/c, etc.). In contrast, the Collaborative Cross (CC) (Threadgill *et al*. 2002; Churchill *et al*. 2004; Collaborative Cross Consortium 2012) is a panel of recombinant inbred (RI) mouse strains, derived from eight founder laboratory strains, that was designed as an optimized murine model of heterogeneous human populations (Churchill *et al*. 2004; Aylor *et al*. 2011; Threadgill *et al*. 2011; Collaborative Cross Consortium 2012). The CC captures the complexity of the mammalian genome, and is an important resource for mapping complex traits and system genetics efforts (Aylor *et al*. 2011; Kelada *et al*. 2012; Ferris *et al*. 2013; Gralinski *et al*. 2015; Gralinski *et al*. 2017; Shorter *et al*. 2017; Srivastava *et al*. 2017). The CC has also been a source of new models of human diseases (Rogala *et al*. 2014; Graham *et al*. 2016; McMullan *et al*. 2018; Orgel *et al*. 2019). By mating different CC strains, we can generate recombinant inbred inter-crosses (CC-RIX) (Zou *et al*. 2005; Schoenrock *et al*. 2018). CC-RIX are outbred but their genomes are completely reproducible (by mating their parental CC strains) and thus are an intriguing model of the genetic diversity akin to that in human populations (Zou *et al*. 2005; Liu *et al*. 2018). The genetic diversity of CC-RIX enables the study of parent-of-origin effects and their phenotypic characterization has demonstrated that there are strong strain effects in some measures (Crowley *et al*. 2015; Schoenrock *et al*. 2018). Using standard inbred lines, we previously demonstrated that we could effectively deliver haloperidol at human-like steady-state concentrations, that steady-state plasma haloperidol concentration is a trait with relatively high heritability, and that there were substantial RI strain differences in the vulnerability to VCMs (Crowley *et al*. 2012a; Crowley *et al*. 2012b; Crowley *et al*. 2014).

In this study, we generated 840 mice and performed extensive phenotypic characterization of 777 mice representing 73 RIX lines chronically treated with haloperidol or placebo in order to gain insight into the genetic basis of ADRs. We performed a battery of behavioral tests, including open field activity, the inclined screen test, and video recordings of VCMs, in order to examine the strain and treatment effects of chronic haloperidol exposure. Using linear mixed models we showed that RIX lines exhibited strong strain effects and strain-by-treatment interactions, supporting the hypothesis of a significant differential contribution of their genetic background to haloperidol-induced ADRs. The fact that RIX lines displayed a range of responses to haloperidol-induced ADRs, with a distribution that is comparable to the susceptibility of haloperidol-ADRs in the human population, supports their use as a model for the study of pharmacogenomics and additional phenotypes.

## MATERIALS AND METHODS

### Mice

A total of 840 mice were tested, including 423 females (210 haloperidol, 213 placebo) and 417 males (210 haloperidol, 207 placebo). These mice represented a total of 73 RIX lines which were derived from 62 CC strains and tested across 51 batches. See ***Table 1*** for summary information on mice with complete data for each phenotype. RIX mice used in this study were born between 5/7/2012 and 6/16/2014. All mice were bred from CC strains from the Systems Genetics Core Facility at the University of North Carolina (http://csbio.unc.edu/CCstatus/index.py?run=availableLines). CC mice were crossed to generate RIX mice as shown in ***Figure 1*** and ***Figure S1***. Pups were weaned at 3 weeks of age and housed two animals per cage, with one randomly assigned to receive haloperidol and the other placebo. Animals were maintained on a 14 hour light/10 hour dark schedule with water and food available *ad libitum*. All testing procedures were conducted in strict compliance with the Guide for the Care and Use of Laboratory Animals (Institute of Laboratory Animal Resources, National Research Council 1996) and approved by the Institutional Animal Care and Use Committee of the University of North Carolina.

**Table 1.** Summary of subjects with complete data for each phenotype.

| Endpoint | # w/ Pre-Treatment Data | # w/ Post Treatment Data | # w/ Complete Data | Placebo |  | Haloperidol |  | RI (parents) | RIX | # Batches |
| --- | --- | --- | --- | --- | --- | --- | --- | --- | --- | --- |
|  |  |  |  | Male | Female | Male | Female |  |  |  |
| Distance | 828 | 785 | 777 | 199 | 204 | 188 | 186 | 62 | 73 | 51 |
| Vertical | 828 | 785 | 777 | 199 | 204 | 188 | 186 | 62 | 73 | 51 |
| Stereotypy | 828 | 785 | 777 | 199 | 204 | 188 | 186 | 62 | 73 | 51 |
| Centroid | 828 | 785 | 777 | 199 | 204 | 188 | 186 | 62 | 73 | 51 |
| VCM | n/a | 731 | 731 | 183 | 192 | 177 | 179 | 62 | 73 | 51 |
| Weight | 823 | 716 | 715 | 179 | 191 | 171 | 174 | 58 | 68 | 46 |
| EPS | 783 | 639 | 604 | 152 | 155 | 150 | 147 | 60 | 68 | 39 |
| Plasma* | n/a | 254 | 254 | - | - | 125 | 129 | 48 | 53 | 37 |

\* applies only to haloperidol arm

**Figure 1.**
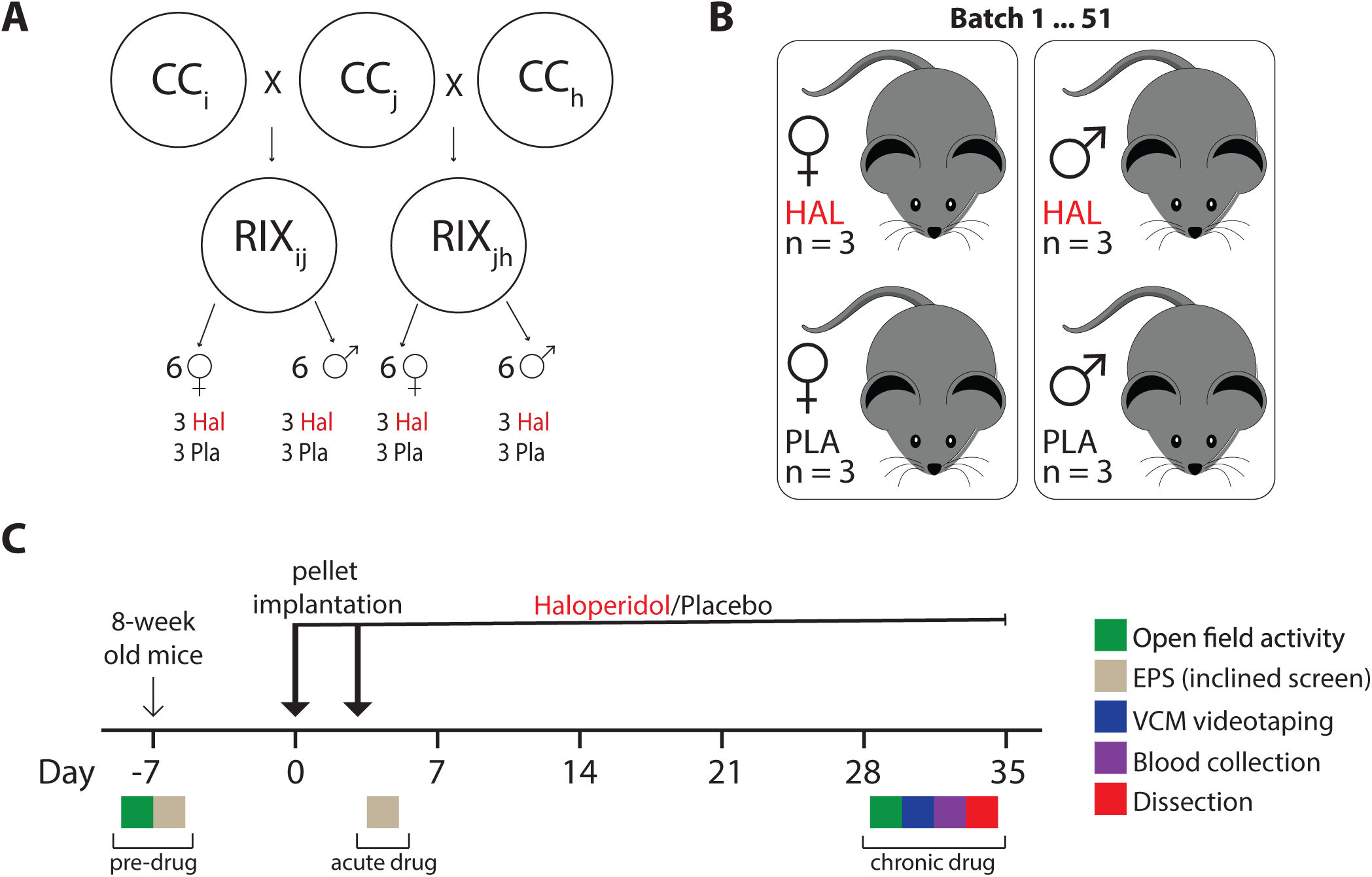
RIX breeding scheme, experimental design and phenotyping pipeline. **(a)** CC mice were crossed in a quasi-loop design (See ***Figure S1***) in order to generate RIX lines with maximum genetic diversity. We tested up to 12 animals per RIX line: 6 males (3 haloperidol, 3 placebo) and 6 females (3 haloperidol, 3 placebo). **(b)** Mice were housed two per cage from weaning with one designated for drug treatment and the other for placebo treatment. To avoid batch effects, each RIX line was tested across up to three batches. **(c)** RIX mice were aged for ∼8 weeks before being added to the phenotyping pipeline. Phenotyping occurred “pre-drug” (7 days before pellet implantation), after “acute drug” treatment (on day 4; one day after split pellet implantation), and chronic treatment with haloperidol or placebo (“chronic drug”: 28-35 days after pellet implantation). Open field activity (OFA) was done both pre-drug and after chronic drug treatment. The inclined screen test, which measures extrapyramidal side effects (EPS) was done both pre-drug and after acute drug treatment. Vacuous chewing movements (VCMs) were measured after chronic drug treatment.

The study design of our 6-week haloperidol phenotyping protocol is summarized in ***Figure 1***. Most RIX lines (63 of 73) included 12 animals, although some lines had fewer (two were composed of only two animals each) and others more (one line included 18 mice). Given the number of mice assayed in this study and the battery of tests we employed, it was unfeasible to generate and test all mice in a single batch. Our mixed models include batch as a random effect to account for this possible confounder.

### Haloperidol exposure

Eight-week old mice (±7 days) were implanted with slow-release haloperidol pellets (3.0 mg/kg/day; Innovative Research of America; Sarasota, FL, USA) (Fleischmann *et al*. 2002) or placebo and treated for 30 days for a chronic haloperidol administration paradigm. All experimental procedures were randomized to minimize batch artifacts (Leek *et al*. 2010) (e.g., assignment to haloperidol or placebo; cage; order of dissection; and assay batch). Experimenters were blind to treatment status.

We have previously demonstrated that this procedure reliably yields human-like steady-state concentrations of haloperidol (typically 3.75–19 ng/ml) (Hsin-Tung and Simpson 2000) in blood plasma and brain tissue. This chronic dosing paradigm reliably results in VCMs (an established model of extrapyramidal symptoms) (Turrone *et al*. 2002b) in multiple mouse strains (Crowley *et al*. 2011; Crowley *et al*. 2012a; Crowley *et al*. 2014). Haloperidol pellets were implanted subcutaneously with a trocar under 2 min of isoflurane anesthesia to minimize handling stress and pain. Two pellets of incremental dosages were implanted 2 days apart to compensate for varying body weights and to minimize acute sedation (Crowley *et al*. 2012a). Placebo-treated animals were implanted with pellets containing the same matrix material but no drug. Following pellet implantation, we positioned food and water near the surface of the bedding, since movement can be compromised in a strain-specific manner due to variable degrees of haloperidol-induced extrapyramidal side effects.

### Open field activity (OFA)

Extrapyramidal side effects may appear as general motor deficits in mice. Open field activity is a behavioral assay that measures general locomotor activity, anxiety, and exploratory drive (Crusio 2013). OFA was measured on days −7 and +28 relative to the start of drug treatment (day 0). Spontaneous locomotor activity in the open field (Crawley 1985) was measured for 30 minutes using a photocell-equipped automated open field apparatus with a white Plexiglas floor and clear Plexiglas walls (Superflex system, Accuscan Instruments, Columbus, OH; arena measures 40 cm wide × 40 cm long × 30 cm high) surrounded by infrared detection beams on the X, Y and Z-axes that track the animals’ position and activity over the course of the experiment. The apparatus is isolated within a 73.5 x 59 x 59 cm testing chamber fitted with overhead fluorescent lighting (lux level 14). One to two hours before testing, cages were moved from the housing room to the testing room. Following acclimation, animals were removed from their home cage, immediately placed in the corner of the open field arena, and allowed to freely explore the apparatus for a 30-minute test interval. Four phenotypes were extracted from these activity data: total distance traveled (cm), vertical activity (number of beam breaks on the Y-axis), stereotypy (a measure of repetitive movements based on repeated breaking of the same beam), and time spent in the central region of the chamber (a measure of anxiety). Activity chambers were located inside of sound-attenuating boxes equipped with houselights and fans.

### Extrapyramidal side effects (EPS)

The inclined screen test (Barnes *et al*. 1990) was used as an index of Parkinsonian rigidity and sedation. Mice were placed on a wire mesh screen inclined at 45° and the latency to move all four paws was recorded (to a maximum of 60 seconds). EPS was measured at baseline (day −5) and 24 hours after implantation of the second drug pellet (day 3), since pilot work indicated that haloperidol-induced EPS was most pronounced after acute, rather than chronic, drug treatment.

### Orofacial movement scoring

Video recording of vacuous chewing movements (VCMs) was carried out after 28 days post-treatment. To this end, mice were briefly anesthetized with isoflurane and restrained for 25 minutes using a plastic collar. Collars were made from two plastic semicircular pieces that were adjusted based on neck size and to achieve the most comfortable position for the mouse. The collar partially immobilized the mice at the neck but still permitted head movement to allow for video recording of jaw movements by JVC Everio digital camcorders. Digital videotapes were made using the protocol developed by Tomiyama et al. (Tomiyama *et al*. 2001). The first 10 minutes of video were not analyzed in order to allow the mice to adjust to the collar and to relax. The last 15 minutes of the video were scored for orofacial movement. Videos were randomized and scored by a single-blinded rater to increase consistency and to reduce any deviation or bias between raters. The rater was trained by an expert and a set of standard training videos used in the studies by Crowley et al. (Crowley *et al*. 2012a; Crowley *et al*. 2012b) to align the rater with the correct identification of VCMs according to the scoring from those previous studies. Drift was monitored by re-scoring random videos throughout the course of the study.

The movements that were specifically analyzed and counted were tongue protrusions, jaw tremors, overt chewing movements, and subtle chewing movements. Individual events of each movement were counted. Subtle chewing movements were defined as instances of vertical jaw movement in which the inside cavity of the mouth could not be seen and the jaw was not open for a long period of time. Overt chewing movements occurred when a larger vertical movement was observed in which the cavity could be seen and the jaw was open for an extended length of time. The videos were scored using The Observer XT (Noldus Inc., Wageningen, Netherlands) observational data analysis program. Previous work by Crowley et al had shown that when 27 inbred strains were treated with chronic haloperidol, the behavioral domains that were measured loaded onto two factors, rather than being discrete constructs (Crowley *et al*. 2012a). One factor loaded primarily on antipsychotic-induced changes in open field activity, while the other loaded primarily on haloperidol-induced orofacial movements. In this study, we have collapsed all four VCM measures into a single factor.

### Tissue collection

Mice were sacrificed on day +32 relative to the start of drug treatment by cervical dislocation. We collected blood plasma from haloperidol-treated mice at this time using EDTA-treated tubes in order to measure haloperidol drug concentration. Blood was centrifuged to isolate plasma. Haloperidol assays were performed using mass spectrometry at the Analytical Psychopharmacology Laboratory located at the Nathan Kline Institute for Psychiatric Research (Orangeburg, NY, USA).

### Statistical models

As shown in ***Table 1***, the phenotypes studied can be grouped into three classes depending on the nature of outcome ascertainment; correspondingly, three separate statistical models were employed. For the open field activity, EPS, and body mass phenotypes, outcomes were ascertained on two occasions (pre- and post-treatment) in both arms. VCMs were measured on a single occasion (post-treatment) in both arms. Blood plasma phenotype was also only measured post-treatment, but naturally only applies to the haloperidol arm.

For the open field, EPS, and body weight phenotypes, the difference between pre- and post-treatment measurements were taken and log-transformed for body weight. The mixed model employed for these phenotypes was:

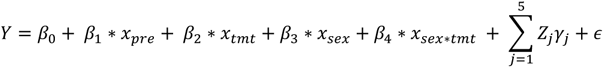

where *Y* refers to the vector of transformed responses, *x*_*pre*_ to the pre-treatment phenotype value, *x*_*tmt*_ is a vector of treatment indicators, *x*_*sex*_ controls for sex, and *x*_*sextmt*_ controls for the interaction of sex and treatment. Each *γ*_*j*_ refers to a random effect vector associated with the *j*th random effect. The *γ*_*j*_′*s* correspond to the batch, strain, strain-by-treatment, strain-by-sex, and strain-by-sex-by-treatment effects (henceforth referred to as effects 1-5, respectively). We assume that the *γ*_*j*_’s are mutually independent and 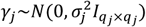. Further, the error term 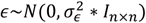, independent of the random effects. Details on the construction of the associated random design matrices are included in ***File S1***.

A single VCM phenotype was derived by summing the number subtle and overt VCM, tongue movements, and tremors. For this phenotype, the following model was employed, with the term to control for the pre-treatment value excluded, as follows:

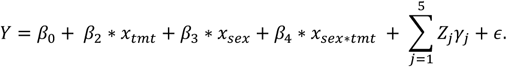

Haloperidol blood concentration was log-transformed, and the following model was employed. This model controls for three random effects: batch, strain, and sex (1, 2, and 4):

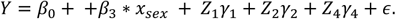

All models were fit in SAS/STAT® 14.1 software, Version 9.4 of the SAS System for Linux (SAS Institute Inc. 2015), using PROC MIXED. This procedure allows the user to specify flexible design matrices, but the required syntax to achieve the desired correlation structure is nontrivial, and sample code is included in ***File S1***.

### Data availability

All data from this study have been deposited in the Mouse Phenome Database (https://phenome.jax.org/). Available CC strains can be obtained from the Systems Genetics Core Facility at the University of North Carolina (http://csbio.unc.edu/CCstatus/index.py?run=availableLines). Supplemental files are available at FigShare. *Table S1* describes the CC strains used to generate the RIX lines that were part of the study. *Figure S1* shows the mating scheme used to cross CC and generate RIX with maximum genetic diversity. *Figure S2* shows haloperidol-induced changes in behavioral measures. *Figure S3* shows the raw data for RIX mice treated with haloperidol or placebo, for the centroid time and stereotypy measures of OFA. *Figure S4* shows RI strain-by-treatment and strain prediction intervals for stereotypy, centroid, and EPS. *Figure S5* presents raw data and strain-by-treatment predictions for body weight of RIX mice treated with haloperidol or placebo. *Figure S6* shows CC strain-level predictions for plasma haloperidol levels. *Figure S7* compares the pre-treatment distributions of various behavioral measures between the CC-RIX, the CC diallel study (Crowley *et al*. 2014), and a survey of 27 mouse strains (Crowley *et al*. 2012a; Crowley *et al*. 2012b). *File S1* includes the sample code.

## RESULTS

### Generation of RIX lines from CC strains

In this study, we used RIX mice derived from the genetically diverse CC in order to understand the effects of haloperidol-induced ADRs, using a battery of behavioral tests as a means of doing a comprehensive phenotypical characterization. To maximize genetic diversity and test outbred mice with reproducible genomes, we generated RIX mice by crossing males and females from two different CC RI strains (***Figure 1a***) in a quasi-loop design (***Figure S1***). This quasi-loop design used all RI genomes available at the time, generating a mostly uniformly structured population, potentially expanding the analytical possibilities beyond simple additive models, and providing a systematic approach to detect regulatory variation including parent-of-origin effects. We performed a battery of behavioral tests in 840 mice from 73 RIX lines (derived from 62 CC strains) treated with haloperidol or placebo in order to monitor the development of ADRs. ***Figure 1c*** details the phenotyping pipeline for this study. Mice were entered into the phenotyping pipeline at 8 weeks of age and behavioral assays were carried out as detailed in *Materials and Methods*. Below, we provide both descriptive as well as model-based results for the behavioral phenotypes examined using the CC-RIX.

### Descriptive Data

#### I. CC-RIX mice exhibit reduced open field activity after chronic treatment with haloperidol

The raw data are visualized at the marginal (that is, not strain-specific) level in a series of density plots (***Figures 2a-d***) that illustrate the densities for each phenotype. Open field activity decreased from pre- to post-treatment for mice treated with placebo or haloperidol (***Figure 2a-b***). This is visualized as a left-shift in the distribution curve, for all phenotypes except for stereotypy (***Figure 2d***). This was expected for both cohorts, as mice are known to spend less time exploring the open field chamber when re-exposed, due to lack of novelty. For distance traveled (***Figure 2a; Figure S2a***) and vertical activity (***Figure 2b; Figure S2b***), the left-shift is far more pronounced for the haloperidol treated mice, suggesting a strong average treatment effect. The left-shift is more modest for time in centroid (***Figure 2c; Figure S2c)***, and not observed at all for stereotypy (***Figure 2d; Figure S2d***). Note that this implies only that there is a lack of a marginal effect. For stereotypy, although the raw data do not suggest a treatment effect at the mean, treatment still affects individual RIX strains differently (see below).

**Figure 2.**
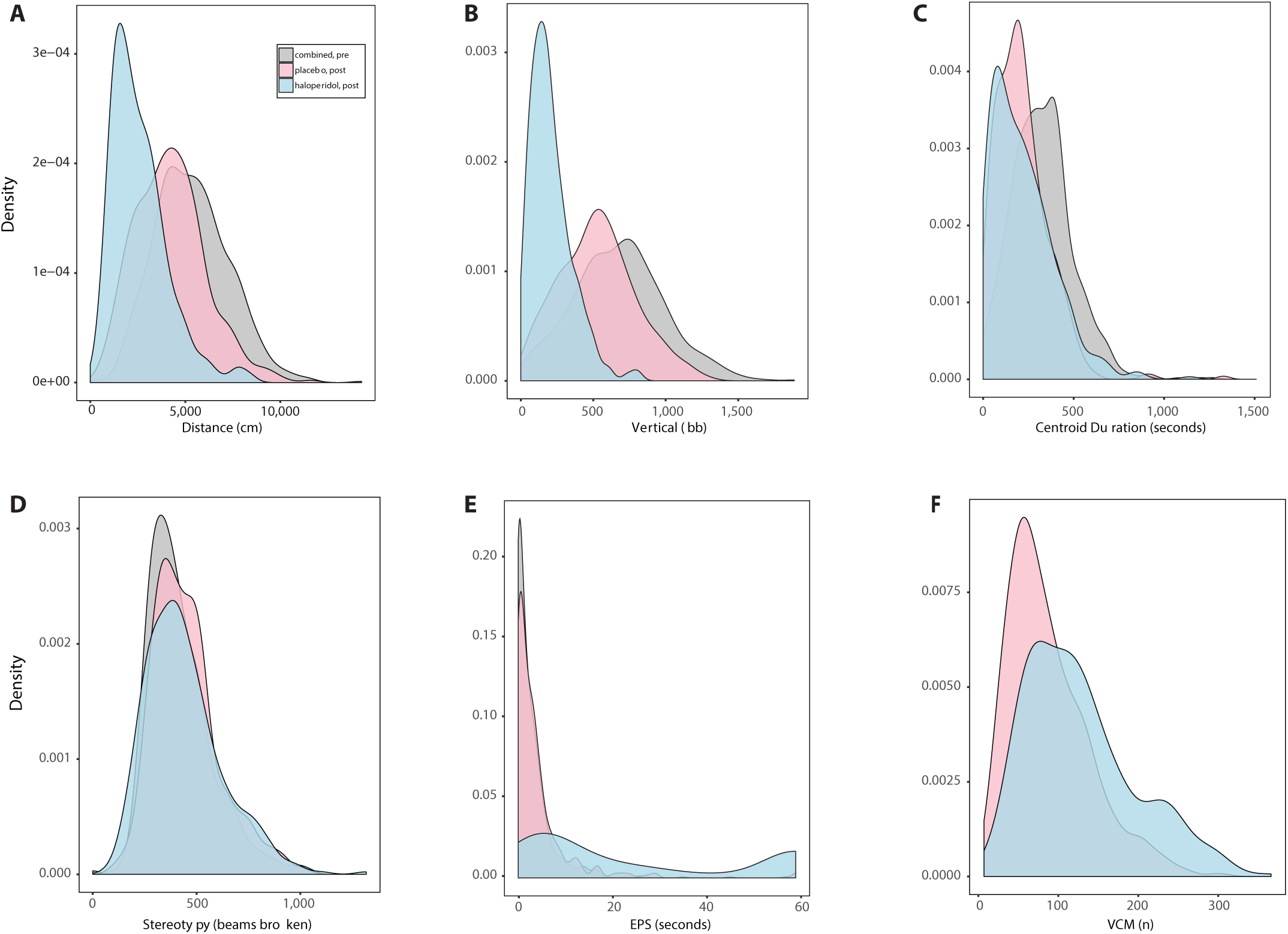
Overlaid series of density plots of individual behavioral measures for haloperidol- and placebo-treated mice, before (pre) and after (post) chronic drug treatment. The gray series is comprised of the combined pre-treatment data for the placebo- and haloperidol-treated samples. Pre-treated data were combined, as haloperidol- and placebo-treated animals were indistinct from each other prior to treatment. The pink and blue series correspond to the post-treatment densities for placebo- and haloperidol-treated samples, respectively. **(a)** Distance traveled (cm) from OFA. **(b)** Vertical activity (beam breaks) from OFA. **(c)** Centroid time (seconds) from OFA. **(d)** Stereotypy (beam breaks) from OFA. **(e)** EPS from inclined screen test (seconds). **(f)** VCMs after chronic treatment with haloperidol or placebo.

#### II. Acute haloperidol exposure results in increased time on inclined screen test

The raw data from the inclined screen test (***Figure 2e***), suggest an extreme effect of haloperidol. For pre-treatment and placebo-treated mice the series cluster around zero; these mice move almost immediately after being placed on the inclined screen. Conversely, the distribution for the haloperidol treated mice is bimodal, exhibiting an extremely fat right tail, enriched with a noticeable proportion of animals who are unable to move their four paws at all within the 60-second experiment.

#### III. Chronic haloperidol treatment increases incidence of VCMs

The raw data for VCM are visualized in the distribution curves in ***Figure 2f***. Haloperidol-treated RIX mice show increased susceptibility to the emergence of VCMs, as evidenced by the fattening of the right tail in the haloperidol series vis-à-vis the placebo series. Note a small but noticeable proportion of animals from both treatment paradigms do not experience any VCMs.

#### IV. Strain-Level Descriptive Figures

***Figures 2a-e*** and ***Figures S2a-e*** provide an overview at the population-average level. ***Figure 3*** and ***Figure S3*** show the raw data at the RIX line level. For each line the distribution of the phenotype (or change in phenotype, where appropriate), is visualized in boxplots where the first and third quartiles are visualized at the bottom and top of the boxes, respectively, and the median as a black hash mark. The pink boxes correspond to the placebo data and the blue to haloperidol. The strains are organized by increasing difference between these strain-level medians. These differences in medians are visualized as the monotonically increasing series of red dots. Chronic haloperidol exposure had divergent effects across OFA measures. For distance traveled and vertical activity, more than half of the RIX lines display a reduction in these OFA measures after haloperidol exposure (***Figure 3a-b***), while less than a quarter of RIX lines displayed changes in stereotypy (***Figure S3a***), and centroid duration (***Figure S3b***) seemed unaffected in most lines. The raw data for EPS showed that the majority of RIX lines showed a delayed response in the inclined screen test after haloperidol treatment (***Figure 3c***). Lastly, the raw data for the VCM measure showed that more than half of the RIX lines displayed higher incidence of VCMs after haloperidol treatment (***Figure 3d***).

**Figure 3.**
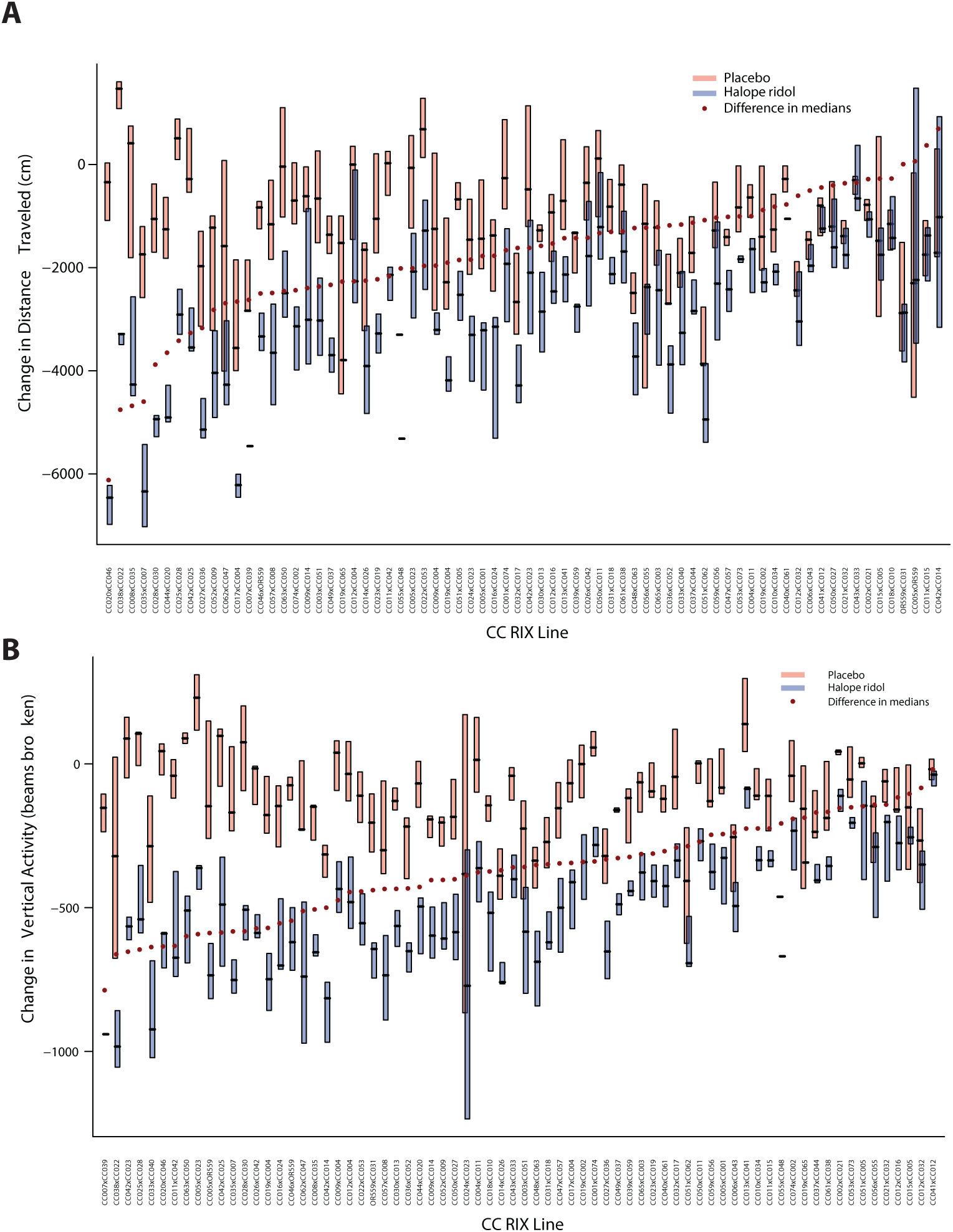
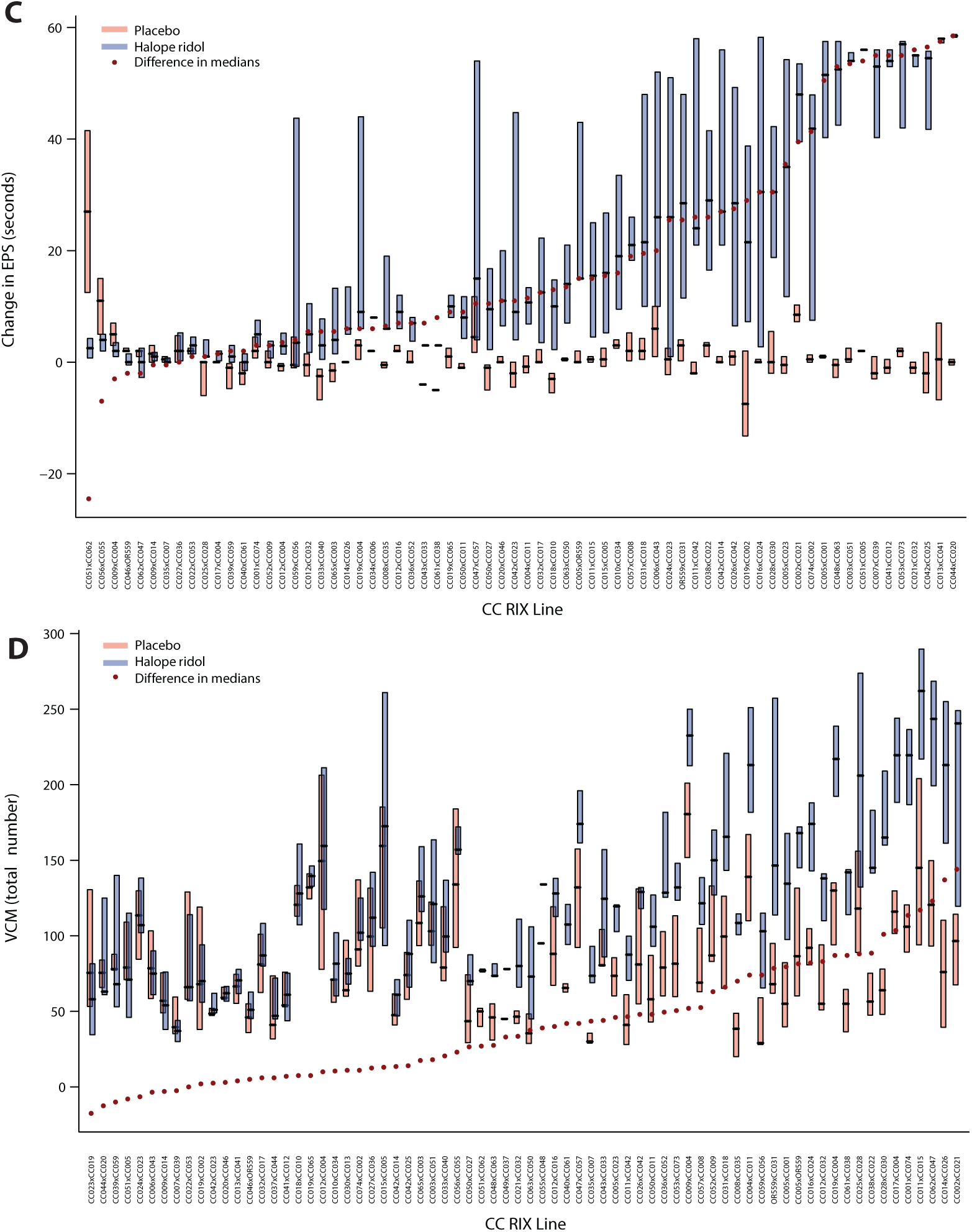
RIX lines exhibited differential sensitivity to haloperidol-induced changes in OFA, EPS, and VCMs. (a) Box plots of individual OFA measures across all phenotyped RIX lines, organized according to their sensitivity to haloperidol-induced changes in distance traveled. (b) Vertical activity. (c) EPS. (d) VCMs

### Model-Fitting Results

#### I. Hypothesis Tests

The main model results are in ***Table 2***, which includes results for all eight phenotypes assayed in our experiment. For ease of readability, some p-values were also not explicitly listed, but rather the level at which they were significant is simply indicated by asterisks and color-coding. The first five rows of the table contain the fixed effect estimates with p-values in parentheses. These p-values were computed using the Kenward-Roger approximation to compute the denominator degrees of freedom. The next four rows contain the estimates for the variance components of interest (the batch effect has been excluded), with the p-values computed from the likelihood ratio tests (LRTs). The next four rows correspond to test statistics and p-values that involve multiple parameters; the numbers in the rows corresponding to fixed effects are F-statistics, and those in the rows corresponding to tests of random effects are likelihood ratio statistics.

**Table 2.** Results from statistical analyses.

| Effect Type | Effect | Estimate (p-value) |  |  |  |  |  |  |  |
| --- | --- | --- | --- | --- | --- | --- | --- | --- | --- |
|  |  | Open Field Phenotypes |  |  |  | VCM | Weight | EPS | Plasma |
|  |  | Distance | Vertical | Centroid | Stereotypy |  |  |  |  |
| Fixed Effects | Intercept | 1354.81** | 150.16** | 71.53** | 287.13** | 105.40** | 0.4860** | 14.743** | 2.5931** |
|  | Pre | -0.65** | -0.69** | -0.70** | -0.65** | -- | -<br>0.1181** | -0.7391** | -- |
|  | Sex | -112.74<br>(0.11) | -0.94<br>(0.91) | 3.23<br>(0.44) | 2.84 (0.73) | 8.0108 * | 0.0092<br>(0.20) | 0.8715<br>(0.29) | 0.03297<br>(0.4329) |
|  | Tmt | -911.69** | -165.80** | 1.27<br>(0.80) | -13.34<br>(0.15) | 18.4336** | 0.0104<br>(0.03) | 10.4421** | -- |
|  | Sex*tmt | 208.26<br>(0.23) | -3.18<br>(0.87) | -8.46<br>(0.48) | 3.38 (0.87) | 4.9340<br>(0.57) | -0.043* | -2.6416<br>(0.26) | -- |
| Variance Components | Strain | 300278** | 7032.48** | 1020.47** | 2599.73** | 611.53** | 8.4e-4** | 19.760** | 0.04127* |
|  | Strain*Tmt | 86343** | 3005** | 114.54<br>(0.013) | 451.66* | 61.24* | 2.9e-5<br>(0.51) | 21.948** | -- |
|  | Strain*Sex | 22771* | 19910<br>(0.04) | 3.72<br>(0.47) | 199.56<br>(0.07) | 3.57<br>(0.39) | 2.1e-4** | 0.00<br>(0.50) | 0.01105* |
|  | Strain*Sex*Tmt | 4618<br>(0.26) | 242 (1.0) | 0.00<br>(0.50) | 47.68 (0.31) | 26.61<br>(0.06) | 0.00<br>(1.0) | 0.1648<br>(0.46) |  |
| Estimates | Fixed Tmt | 50.50** | 90.44** | 0.27<br>(0.76) | 1.39 (0.25) | 27.08** | 5.358* | 25.99** | -- |
|  | Fixed Sex | 1.32<br>(0.27) | 0.05<br>(0.96) | 0.33<br>(0.72) | 0.18 (0.84) | 13.20** | 5.37* | 0.72<br>(0.4894) |  |
|  | Random Tmt | 46.09** | 74.30** | 4.91<br>(0.03) | 8.347* | 10.58* | 0.29<br>(0.51) | 40.82** | -- |
|  | Random Sex | 6.78<br>(0.013) | 3.15<br>(0.09) | 0.006<br>(0.72) | 2.368<br>(0.139) | 2.45<br>(0.13) | 9.95* | 0.012<br>(0.71) |  |
| Overall Heritability |  | 0.4 | 0.33 | 0.24 | 0.24 | 0.40 | 0.21 | 0.28 | 0.31 |
| Treatment effect heritability |  | 0.09 | 0.12 | 0.02 | 0.04 | 0.05 | 0.01 | 0.15 | -- |

\* $P < 0.01$ (light grey)\*\* $P < 0.001$ (dark grey)

One salient result is the strong marginal (fixed) effect of treatment, which is highly significant for most phenotypes. Confirming the results from the exploratory analyses above, it is not significant in centroid and stereotypy models, however, and only weakly significant in the model for body weight. Note, however, that the strain effect is highly significant for all models, including centroid and stereotypy. The random treatment effects are also significant for all phenotypes except body weight (***Figure S5***).

#### II. Best linear unbiased predictions (BLUPS) to visualize CC strain level predictions

The main model results included in ***Table 2*** provide evidence of the existence of strain and strain-by-treatment interactions. However, determining which parental (RI) strains contribute most is also of interest; i.e., we are interested in the levels of the random effects not just the estimates of the variance components. We used BLUPS for these strain level predictions. We plotted these in ascending order for all phenotypes (***Figure 4***). The absolute height of each bar corresponds to the prediction interval. Due to the small number of samples per RIX, there is little power to differentiate between the RI effects: in comparing two strain predictions, their intervals tend to overlap, except when comparing strains at the extremes. Under the assumption of the additive strain effects, these strain-level predictions can be used to predict which RIX lines will exhibit extreme phenotypes.

**Figure 4.**
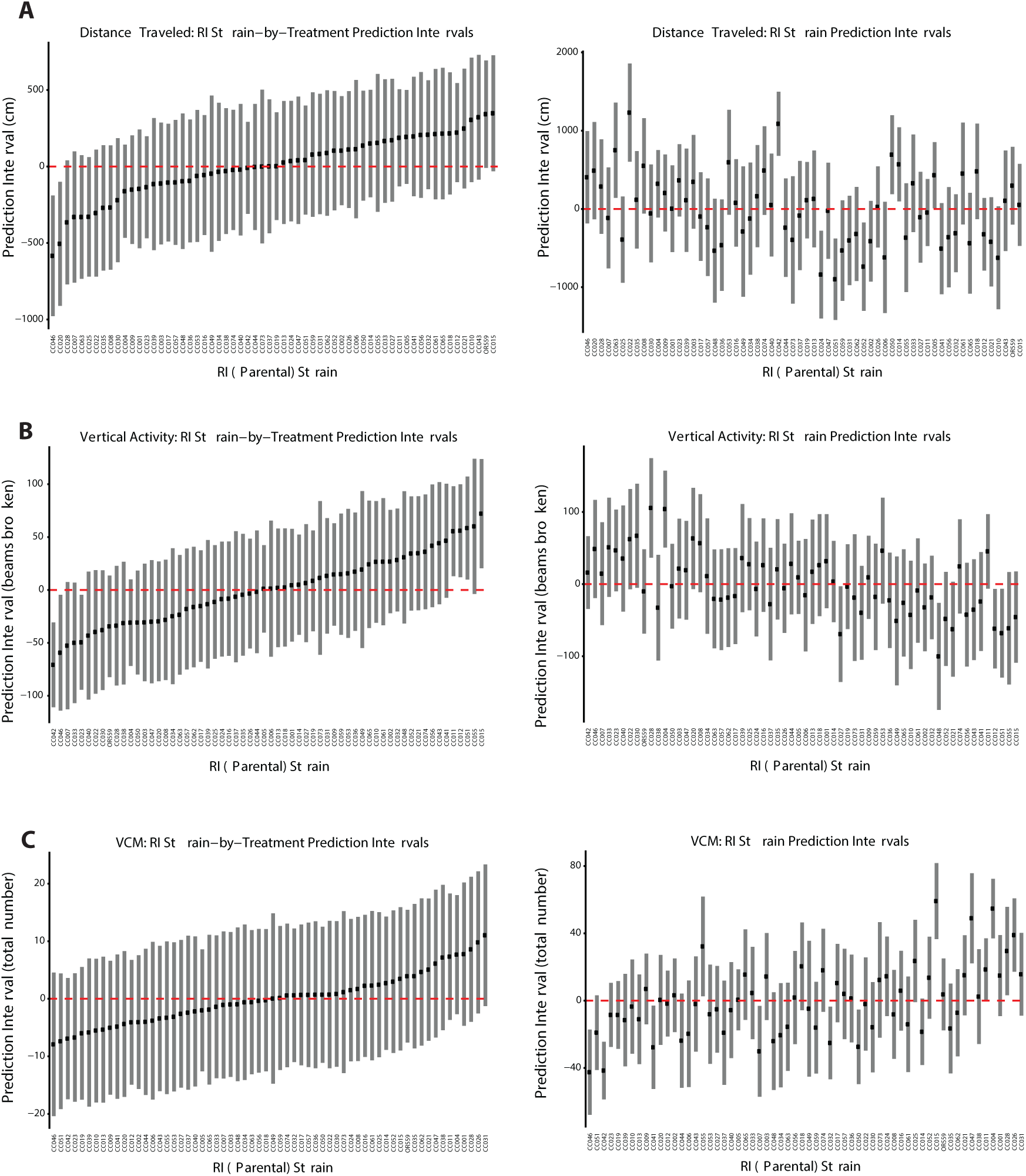
RI strain-by-treatment (left) and strain (right) prediction intervals for: (a) Distance traveled, (b) vertical activity, and (c) VCMs.

#### III. Strong correlations across phenotypes and behavioral measures

We can leverage information across phenotypes to find strains that have consistently extreme predictions, and use this information to actually inform future experiments by selecting strains that have the most extreme results. Here, we show that the strain predictions are correlated across phenotypes, as are strain-by-treatment interactions. To assess this correlation, we used the Pearson correlation coefficient and present the results in the correlograms in ***Figure 5***. The first panel applies to the strain predictions and the second to the strain-by-treatment interactions (note that the blood haloperidol concentration appears only in this second panel; although the relevant random effects were termed “strain” effects in the model for this phenotype, they were only predicted in haloperidol-treated mice). For the additive strain effects, there are two salient results. The distance, vertical and body weight phenotypes are correlated, with changes in horizontal distance correlating highly with changes in vertical beams broken. Further, those strains with the largest decreases in the phenotypes exhibited the smallest increases in weight, as expected. The other two open field phenotypes exhibit some correlation with VCM. Increased time spent in centroid is associated with fewer VCM, while increased stereotypy is associated with increased VCM. The correlation manifests in the strain-by-treatment effect as well, with the effect of treatment on changes in distance, vertical, and centroid effects all being correlated. Further, the effect of treatment on the changes in distance and time spent in centroid are inversely proportional to the effect of treatment on body weight.

**Figure 5.**
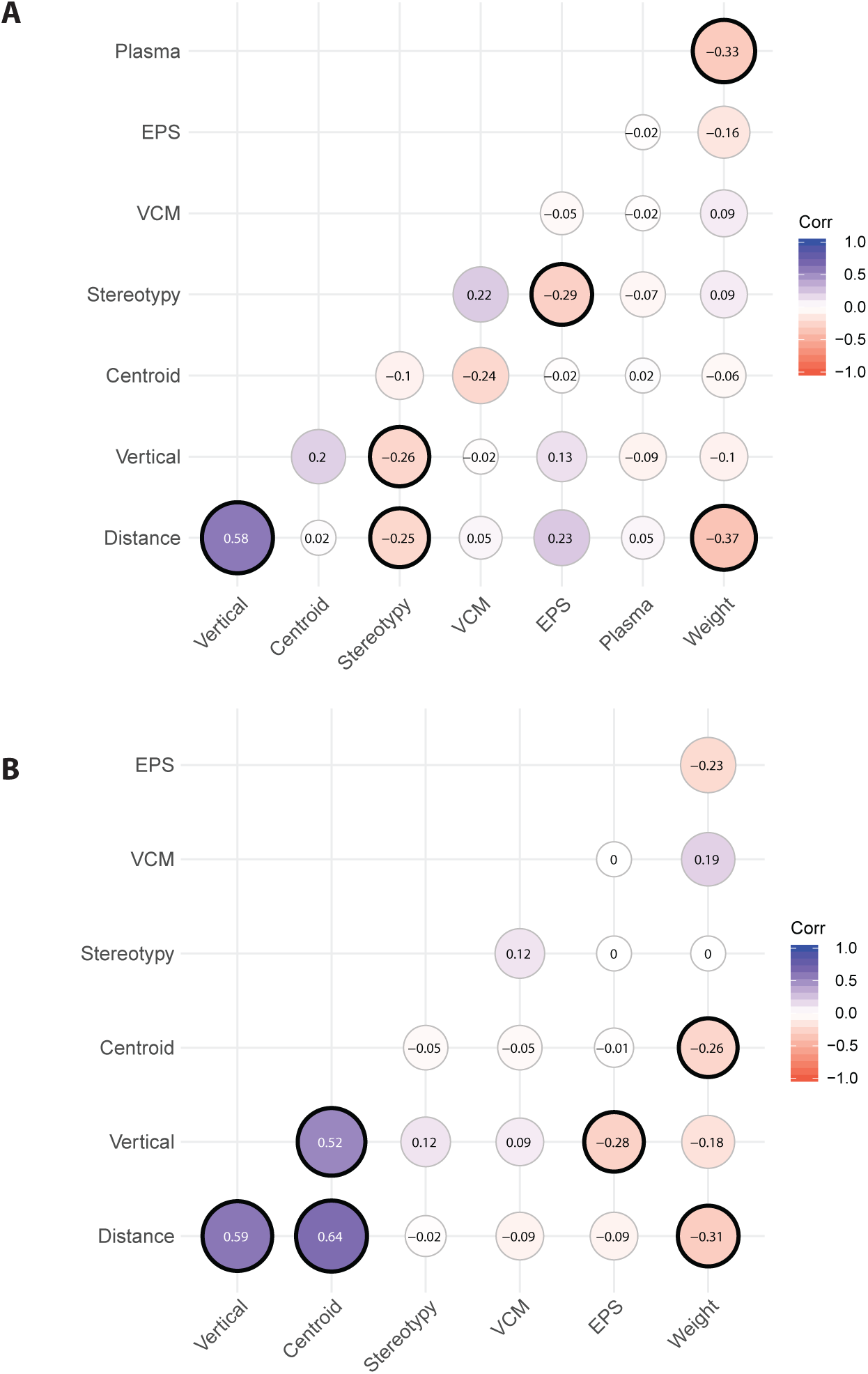
Correlations across phenotypes and behavioral measures for (a) strain and (b) strain-by-treatment effects.

## DISCUSSION

Our study employed the genetically diverse CC-RIX mice to elucidate the genetic basis of antipsychotic ADRs. We performed a battery of behavioral tests in 840 mice from 73 RIX lines treated with haloperidol or placebo in order to monitor the development of ADRs. CC-RIX lines displayed a wide-ranging response to haloperidol treatment induced ADRs as captured in pre- and post-drug behavioral measures. On average, treatment had a strong effect for every phenotype, with the exceptions of centroid and stereotypy. However, note that even for these phenotypes, there was significant between-strain variability. The majority of the CC-RIX lines exhibited reduced distance traveled and vertical activity upon haloperidol treatment, with about a third of the RIX lines exhibiting little change in these measures after exposure to haloperidol (***Figure 3a-b***). Centroid time and stereotypy measures did not appear to be as sensitive to haloperidol induced response (***Figure S3***). The majority of RIX lines showed a delayed response in the inclined screen test (EPS) after haloperidol treatment (***Figure 3c***) and more than half of the RIX lines displayed higher incidence of VCMs after haloperidol treatment (***Figure 3d***). Interestingly, the prevalence of ADRs in the CC-RIX population is analogous to what has been observed in humans, where 20-43% of patients treated with antipsychotics can develop tardive dyskinesia (Halliday *et al*. 2002; Soares-Weiser and Fernandez 2007)

We used linear mixed models to test for the existence of strain and treatment effects. Of note, all phenotype models that included a pre-treatment fixed effect demonstrated significant regression to the mean; subjects with higher pre-treatment scores experienced larger reductions in the phenotype. We observed highly significant strain effects for almost all behavioral measurements investigated (p<0.001). Further, we observed strong strain-by-treatment interactions for most phenotypes, with particularly strong effects (p<0.001) for change in distance traveled and change in vertical activity, as well as change in EPS. When viewing the BLUPs, it is important to keep in mind that given the small sample sizes for each parental strain, comparisons between the effects of individual pairs of parental strains are not necessarily significant, even when the overall effects are highly significant. To draw closer attention to the treatment effect BLUPs, we indicate strains that are at the extreme for those phenotypes with a substantial mean shift in ***Figure 3*** and ***Figure S3*** (in the direction consistent with this mean shift) in the rightmost columns of ***Table S1***. We flag strains that are in the most extreme 10%, finding that 27 strains had extreme responses for at least one of these phenotypes, with eight of these being extreme for multiple phenotypes.

Despite extensive efforts, we have a limited understanding of the heritability of motoric ADRs in humans (Zhang and Malhotra 2011; Drago *et al*. 2012; Crowley *et al*. 2014). As shown in ***Table 2***, estimates of heritability for the phenotypes measured using the CC-RIX ranged from 0.21 (for change in body weight) to 0.4 (for both number of VCMs and change in distance traveled). Using a similar chronic haloperidol treatment paradigm in 27 genetically diverse inbred strains, we previously showed that haloperidol-induced ADRs (including VCMs and EPS) are highly heritable (∼0.9), with strain being a major predictor of phenotypic variation independently of haloperidol plasma levels (Crowley *et al*. 2012a). The discrepancy between the heritability estimates found for VCMs between these studies is likely due to the genetic diversity present in the CC-RIX, when compared to the 27 inbred lines, and the lower heritability found in the CC-RIX (*h*^*2*^∼ 0.4), which is likely closer to the heritability of the analogous trait (TD) in humans.

One major goal of our study is to identify specific genes driving antipsychotic ADRs in the CC-RIX mice. For each open field phenotype, we attempted genetic mapping using R/qtl2. (Broman *et al*. 2019) We did this in two ways, first using pre-treatment phenotype as the dependent variable (controlling for sex) and also using change from baseline as the dependent variable (controlling for pre-treatment phenotype, sex, treatment, and the interaction thereof). We did not detect any LOD peaks that exceeded the significance threshold established based on 1000 permutations. The inability to find significant associations is not unexpected, as these traits appear to be polygenic and we would likely need a much larger sample size to achieve enough power to map these. We have also explored the genetic architecture of these behavioral measures using a diallel cross of the eight genetically diverse founder strains of the CC, and shown that ∼70% of the variance in EPS is explained by parent-of-origin and additive effects (Crowley *et al*. 2014). We compared the pre-treatment distributions of the phenotypes for which comparable data were available in our data, this diallel cross, and the aforementioned survey of 27 mouse strains (see ***Figure S7***). The distributions largely accord with our expectations; for the distance and vertical phenotypes, the right tails of the RIX distributions are thicker than those for the other samples, consistent with their expected hybrid vigor. The same pattern is observed for VCM. Conversely, the mice in the Strain Survey dataset are the heaviest, as they are enriched with NZO samples, which were specifically bred to express this phenotype.

Our study supports the use of the CC-RIX mouse population for pharmacogenomics studies. We observed that 30% of CC-RIX mice exhibited a sensitivity to haloperidol exposure, evidenced by VCMs and EPS, closely matching the proportion of humans that develop TD after treatment with this antipsychotic. Given the significant impact that antipsychotic ADRs and subsequent discontinuation of treatment can have on patients, making headway in our understanding of the genetic basis for the susceptibility to antipsychotic ADRs could advance the development of safer and more effective therapeutic approaches for the treatment of schizophrenia.

## ACKNOWLEDGEMENTS

Major funding was provided by National Institute of Mental Health/National Human Genome Research Institute Center of Excellence for Genome Sciences grants (P50MH090338 and P50HG006582. The Collaborative Cross project is also supported by the University Cancer Research Funds granted to the Lineberger Comprehensive Cancer Center (MCR012CCRI). We thank the High-Throughput Sequencing Facility and the Systems Genetics Core at UNC for their assistance with this project.

## REFERENCES

American Psychiatric Association., 2013 Diagnostic and statistical manual of mental disorders : DSM-5. American Psychiatric Association, Washington, D.C.

Aylor, D. L., W. Valdar, W. Foulds-Mathes, R. J. Buus, R. A. Verdugo et al., 2011 Genetic analysis of complex traits in the emerging Collaborative Cross. Genome Res 21: 1213–1222.

Bakker, P. R., P. N. van Harten and J. van Os, 2006 Antipsychotic-induced tardive dyskinesia and the Ser9Gly polymorphism in the DRD3 gene: a meta analysis. Schizophrenia research 83: 185–192.

Barnes, D. E., B. Robinson, J. G. Csernansky and E. P. Bellows, 1990 Sensitization versus tolerance to haloperidol-induced catalepsy: multiple determinants. Pharmacol Biochem Behav 36: 883–887.

Broman, K. W., D. M. Gatti, P. Simecek, N. A. Furlotte, P. Prins et al., 2019 R/qtl2: Software for Mapping Quantitative Trait Loci with High-Dimensional Data and Multiparent Populations. Genetics 211: 495–502.

Churchill, G. A., D. C. Airey, H. Allayee, J. M. Angel, A. D. Attie et al., 2004 The Collaborative Cross, a community resource for the genetic analysis of complex traits. Nat Genet 36: 1133–1137.

Cloutier, M., M. S. Aigbogun, A. Guerin, R. Nitulescu, A. V. Ramanakumar et al., 2016 The Economic Burden of Schizophrenia in the United States in 2013. J Clin Psychiatry 77: 764–771.

Collaborative Cross Consortium, 2012 The genome architecture of the Collaborative Cross mouse genetic reference population. Genetics 190: 389–401.

Crawley, J. N., 1985 Exploratory behavior models of anxiety in mice. Neurosci Biobehav Rev 9: 37–44.

Crowley, J., Y. Kim, J. Szatkiewicz, A. Pratt, C. Quackenbush et al., 2011 Genome-wide association mapping of loci for antipsychotic-induced extrapyramidal symptoms in mice. Mammalian Genome 23: 322–335.

Crowley, J. J., D. E. Adkins, A. L. Pratt, C. R. Quackenbush, E. J. van den Oord et al., 2012a Antipsychotic-induced vacuous chewing movements and extrapyramidal side effects are highly heritable in mice. Pharmacogenomics J 12: 147–155.

Crowley, J. J., Y. Kim, A. B. Lenarcic, C. R. Quackenbush, C. J. Barrick et al., 2014 Genetics of adverse reactions to haloperidol in a mouse diallel: a drug-placebo experiment and Bayesian causal analysis. Genetics 196: 321–347.

Crowley, J. J., Y. Kim, J. P. Szatkiewicz, A. L. Pratt, C. R. Quackenbush et al., 2012b Genome-wide association mapping of loci for antipsychotic-induced extrapyramidal symptoms in mice. Mamm Genome 23: 322–335.

Crowley, J. J., V. Zhabotynsky, W. Sun, S. Huang, I. K. Pakatci et al., 2015 Analyses of allele-specific gene expression in highly divergent mouse crosses identifies pervasive allelic imbalance. Nat Genet 47: 353–360.

Crusio, W. E., 2013 The genetics of exploratory behavior, pp. 148–154 in Behavioral Genetics of the Mouse: Volume 1: Genetics of Behavioral Phenotypes, edited by F. Sluyter, R. T. Gerlai, S. Pietropaolo and W. E. Crusio. Cambridge University Press, Cambridge.

Drago, A., I. Giegling, M. Schafer, A. M. Hartmann, H. J. Moller et al., 2012 No association of a set of candidate genes on haloperidol side effects. PLoS One 7: e44853.

Ferris, M. T., D. L. Aylor, D. Bottomly, A. C. Whitmore, L. D. Aicher et al., 2013 Modeling host genetic regulation of influenza pathogenesis in the collaborative cross. PLoS Pathog 9: e1003196.

Fleischmann, N., G. Christ, T. Sclafani and A. Melman, 2002 The effect of ovariectomy and long-term estrogen replacement on bladder structure and function in the rat. J Urol 168: 1265–1268.

Graham, J. B., J. L. Swarts, C. Wilkins, S. Thomas, R. Green et al., 2016 A Mouse Model of Chronic West Nile Virus Disease. PLoS Pathog 12: e1005996.

Gralinski, L. E., M. T. Ferris, D. L. Aylor, A. C. Whitmore, R. Green et al., 2015 Genome Wide Identification of SARS-CoV Susceptibility Loci Using the Collaborative Cross. PLoS Genet 11: e1005504.

Gralinski, L. E., V. D. Menachery, A. P. Morgan, A. L. Totura, A. Beall et al., 2017 Allelic Variation in the Toll-Like Receptor Adaptor Protein Ticam2 Contributes to SARS-Coronavirus Pathogenesis in Mice. G3 (Bethesda) 7: 1653–1663.

Halliday, J., S. Farrington, S. Macdonald, T. MacEwan, V. Sharkey et al., 2002 Nithsdale Schizophrenia Surveys 23: movement disorders. 20-year review. Br J Psychiatry 181: 422–427.

Hsin-Tung, E., and G. Simpson, 2000 Medication-induced movement disorders in Comprehensive Textbook of Psychiatry, edited by H. I. Kaplan and B. J. Sadock. Lippincott, Williams and Wilkins, Philadephia, PA.

Kelada, S. N., D. L. Aylor, B. C. Peck, J. F. Ryan, U. Tavarez et al., 2012 Genetic analysis of hematological parameters in incipient lines of the collaborative cross. G3 (Bethesda) 2: 157–165.

Kim, Y., P. Giusti-Rodriguez, J. J. Crowley, J. Bryois, R. J. Nonneman et al., 2018 Comparative genomic evidence for the involvement of schizophrenia risk genes in antipsychotic effects. Mol Psychiatry 23: 708–712.

Laursen, T. M., T. Munk-Olsen and M. Vestergaard, 2012 Life expectancy and cardiovascular mortality in persons with schizophrenia. Curr Opin Psychiatry 25: 83–88.

Leek, J. T., R. B. Scharpf, H. C. Bravo, D. Simcha, B. Langmead et al., 2010 Tackling the widespread and critical impact of batch effects in high-throughput data. Nature reviews. Genetics 11: 733–739.

Lerer, B., R. H. Segman, E. C. Tan, V. S. Basile, R. Cavallaro et al., 2005 Combined analysis of 635 patients confirms an age-related association of the serotonin 2A receptor gene with tardive dyskinesia and specificity for the non-orofacial subtype. The international journal of neuropsychopharmacology / official scientific journal of the Collegium Internationale Neuropsychopharmacologicum (CINP) 8: 411–425.

Lieberman, J. A., T. S. Stroup, J. P. McEvoy, M. S. Swartz, R. A. Rosenheck et al., 2005 Effectiveness of antipsychotic drugs in patients with chronic schizophrenia. N Engl J Med 353: 1209–1223.

Liu, Y., S. Xiong, W. Sun and F. Zou, 2018 Joint Analysis of Strain and Parent-of-Origin Effects for Recombinant Inbred Intercrosses Generated from Multiparent Populations with the Collaborative Cross as an Example. G3 (Bethesda) 8: 599–605.

McMullan, R. C., M. T. Ferris, T. A. Bell, V. D. Menachery, R. S. Baric et al., 2018 CC002/Unc females are mouse models of exercise-induced paradoxical fat response. Physiol Rep 6: e13716.

Miller, D. D., S. N. Caroff, S. M. Davis, R. A. Rosenheck, J. P. McEvoy et al., 2008 Extrapyramidal side-effects of antipsychotics in a randomised trial. Br J Psychiatry 193: 279–288.

Orgel, K., J. M. Smeekens, P. Ye, L. Fotsch, R. Guo et al., 2019 Genetic diversity between mouse strains allows identification of the CC027/GeniUnc strain as an orally reactive model of peanut allergy. J Allergy Clin Immunol 143: 1027–1037 e1027.

Patsopoulos, N. A., E. E. Ntzani, E. Zintzaras and J. P. Ioannidis, 2005 CYP2D6 polymorphisms and the risk of tardive dyskinesia in schizophrenia: a meta-analysis. Pharmacogenetics and genomics 15: 151–158.

Rogala, A. R., A. P. Morgan, A. M. Christensen, T. J. Gooch, T. A. Bell et al., 2014 The Collaborative Cross as a resource for modeling human disease: CC011/Unc, a new mouse model for spontaneous colitis. Mamm Genome 25: 95–108.

Saha, S., D. Chant and J. McGrath, 2007 A systematic review of mortality in schizophrenia: is the differential mortality gap worsening over time? Archives of general psychiatry 64: 1123–1131.

SAS Institute Inc., 2015 SAS/STAT® 14.1 User’s Guide.. Cary, NC: SAS Institute Inc.

Schoenrock, S. A., D. Oreper, J. Farrington, R. C. McMullan, R. Ervin et al., 2018 Perinatal nutrition interacts with genetic background to alter behavior in a parent-of-origin-dependent manner in adult Collaborative Cross mice. Genes Brain Behav 17: e12438.

Shorter, J. R., F. Odet, D. L. Aylor, W. Pan, C. Y. Kao et al., 2017 Male Infertility Is Responsible for Nearly Half of the Extinction Observed in the Mouse Collaborative Cross. Genetics 206: 557–572.

Soares-Weiser, K., and H. H. Fernandez, 2007 Tardive dyskinesia. Seminars in neurology 27: 159–169.

Srivastava, A., A. P. Morgan, M. L. Najarian, V. K. Sarsani, J. S. Sigmon et al., 2017 Genomes of the Mouse Collaborative Cross. Genetics 206: 537–556.

Threadgill, D. W., K. W. Hunter and R. W. Williams, 2002 Genetic dissection of complex and quantitative traits: from fantasy to reality via a community effort. Mamm Genome 13: 175–178.

Threadgill, D. W., D. R. Miller, G. A. Churchill and F. P. de Villena, 2011 The collaborative cross: a recombinant inbred mouse population for the systems genetic era. ILAR J 52: 24–31.

Tomiyama, K., F. N. McNamara, J. J. Clifford, A. Kinsella, N. Koshikawa et al., 2001 Topographical assessment and pharmacological characterization of orofacial movements in mice: dopamine D(1)-like vs. D(2)-like receptor regulation. Eur J Pharmacol 418: 47–54.

Turrone, P., G. Remington and J. N. Nobrega, 2002a The vacuous chewing movement (VCM) model of tardive dyskinesia revisited: is there a relationship to dopamine D(2) receptor occupancy? Neuroscience and biobehavioral reviews 26: 361–380.

Turrone, P., G. Remington and J. N. Nobrega, 2002b The vacuous chewing movement (VCM) model of tardive dyskinesia revisited: is there a relationship to dopamine D(2) receptor occupancy? Neurosci Biobehav Rev 26: 361–380.

World Health Organization, 2008 The Global Burden of Disease: 2004 Update. WHO Press, Geneva.

Zhang, J. P., and A. K. Malhotra, 2011 Pharmacogenetics and antipsychotics: therapeutic efficacy and side effects prediction. Expert Opin Drug Metab Toxicol 7: 9–37.

Zou, F., J. A. Gelfond, D. C. Airey, L. Lu, K. F. Manly et al., 2005 Quantitative trait locus analysis using recombinant inbred intercrosses: theoretical and empirical considerations. Genetics 170: 1299–1311.

